# Expansion of cortical layers 2 and 3 in human left temporal cortex associates with verbal intelligence

**DOI:** 10.1101/2021.02.07.430103

**Authors:** DB Heyer, R Wilbers, AA Galakhova, E Hartsema, S Braak, S Hunt, MB Verhoog, ML Muijtjens, EJ Mertens, S Idema, JC Baayen, P de Witt Hamer, M Klein, M McGraw, ES Lein, CPJ de Kock, HD Mansvelder, NA Goriounova

## Abstract

The expansion of supragranular cortical layers is thought to have enabled evolutionary development of human cognition and language. However, whether increased volume of supragranular cortical layers can actually support greater cognitive and language abilities in humans has not been demonstrated. Here, we find that subjects with higher general and verbal intelligence test (VIQ) scores have selectively expanded layers 2 and 3 only in the left temporal cortex, an area associated with language and IQ-test performance. This expansion is accompanied by lower neuron densities and larger cell-body size. Furthermore, individuals with higher VIQ scores had neurons with larger dendritic trees in left temporal cortex, potentially impacting their function. Indeed, neurons of subjects with higher VIQ scores had faster action potential upstroke kinetics, which improves information processing. These data show that expansion of supragranular layer volume, cortical and cellular micro-architecture and function are associated with improved verbal mental ability in human subjects.

## Introduction

Higher-order functions of the human brain, such as reasoning and language, rely on the neocortex. Cortical expansion and increased neuronal complexity are regarded as the substrates for the evolutionary development of higher cognitive functions that distinguish humans from other species ^1,2^. In particular, supragranular cortical layers 2 and 3 (L2/L3) in humans have a disproportionally high volume compared to primates ^3^, and hold principal neurons with extensive dendritic trees and rare functional properties optimized for information processing ^4–8^. Far from passively integrating the incoming inputs, the different branches of the dendritic tree act as separate computational elements, so that a single neuron is equivalent in computational power to a multi-layer neural network made up of several nodes ^9–11^. Indeed, human neurons in supragranular cortical layers have 3-fold larger and more complex dendritic trees than other species ^5^ and human dendrites were recently shown to perform combinations of logical operations similar to a multi-layered network ^12^. Larger human neurons also have improved input-output performance, transferring synaptic input to action potential (AP) output with higher bandwidth ^4^, as APs in neurons with larger dendrites have faster upstrokes ^7,8^. Whether such neuronal properties can support verbal cognitive ability in humans has not been tested.

In healthy subjects, total cortical thickness associates with full-scale IQ test scores ^13–15^. However, whether this is explained by selective expansion of supragranular layers 2 and 3 is not known due to insufficient resolution of brain-imaging. Furthermore, to understand the impact of increased cortical layer volume and cell size for cognitive brain function, a direct comparison to other primate species is inherently problematic due to profound differences in cognitive behaviour and learning ^16^. This is even more so the case in regards to the cornerstone of human cognition: the ability to use language ^17^. In the human brain, neural substrates of the language system are distributed over cortical areas in temporal, frontal and parietal lobes and are lateralized to the left hemisphere in 96% of the population ^18^. This left lateralisation offers an opportunity to investigate the association between verbal intelligence and cortical architecture and cellular parameters, since these associations must be limited to the left hemisphere.

Here we tested whether verbal cognitive abilities associate with cortical microstructure by collecting temporal cortical tissue from 59 subjects undergoing neurosurgical treatment of predominantly epilepsy or tumors (Supplementary Table 1). The tissue originated exclusively from middle temporal gyrus (MTG, Brodmann area 21) from the left or right hemisphere. Although the resected MTG cortex is not essential for speech (is not part of classical Broca and Wernicke areas), several lines of evidence point to this region as an important site in verbal cognition. Analysis of localized lesions, functional imaging and positron emission tomography in large cohorts of subjects identified this region as a part of the semantic system serving concept and word retrieval and categorization^19 20^. Furthermore, multiple studies of patients with lesions in MTG show that this area is strongly associated with language comprehension and specific semantic deficits ^21–24^. In addition, recordings of single neuron activity in awake neurosurgery patients performing verbal tasks show that MTG is the area of temporal cortex where neurons selectively respond to language and verbal memory tasks ^25,26^. Because of this role of the left MTG as an integral part of the semantic processing network that underlies verbal cognition, we collected Verbal IQ scores (VIQ) as well as Full Scale (FSIQ) and Performance IQ scores (PIQ) from Wechsler Adult Intelligence Scale (WAIS IV) tests that subjects underwent shortly before surgery.

Here we show that individual differences in verbal intelligence in human subjects can be explained by differences in cortical and cellular architecture and function. Our findings provide evidence supporting the notion that biological factors of evolutionary brain development can be driving factors for emerging human mental abilities.

## Results

### Subjects with higher general and verbal IQ scores have thicker cortical thickness in the left MTG due to the selective expansion of L2/L3

Verbal intelligence has been shown to have strong structural correlates in the brain, including a prominent increase in cortical thickness exclusively in the left temporal lobe of subjects with higher VIQ scores ^14^. However, it is not known whether this expansion is due to the upscaling of all or only specific cortical layers. To investigate this we collected neurosurgically resected left or right MTG from 59 subjects treated for epilepsy or tumor (Supplementary table 1). We first asked whether dimensions of specific cortical layers in left or right MTG associate with IQ scores. To visualize cortical layer boundaries, we DAPI stained cortical sections and quantified the thickness of the cortical layers at 4 to 5 locations from multiple (range = 1-9, median = 2) slices per subject (Fig 1a, b). To minimize biological variation, we quantified layer thickness only at the crown of the gyrus (Fig 1b). We calculated average layer thickness per subject and compared 2 groups of subjects with low and high verbal IQ (VIQ) scores. Only in the left and not right MTG, we found that subjects with high VIQ scores (VIQ>90) have selectively expanded cortical L2/L3 compared to subjects with lower VIQ scores (VIQ<90), while other layers were similar across the VIQ groups (Fig. 1c). We also observed a strong positive correlation between individual VIQ scores of the subjects and total cortical thickness and L2/L3 thickness in the left MTG (R=0.8, R^2^ = 0.66; Fig. 1d, Supplementary Fig 1, Fig. 1d). In contrast, VIQ scores did not correlate with the thickness of other cortical layers (Supplementary Fig. 1), suggesting that the expansion of L2/L3 in subjects with higher verbal intelligence underlies the increase in total cortical thickness of the left MTG.

**Figure 1.**
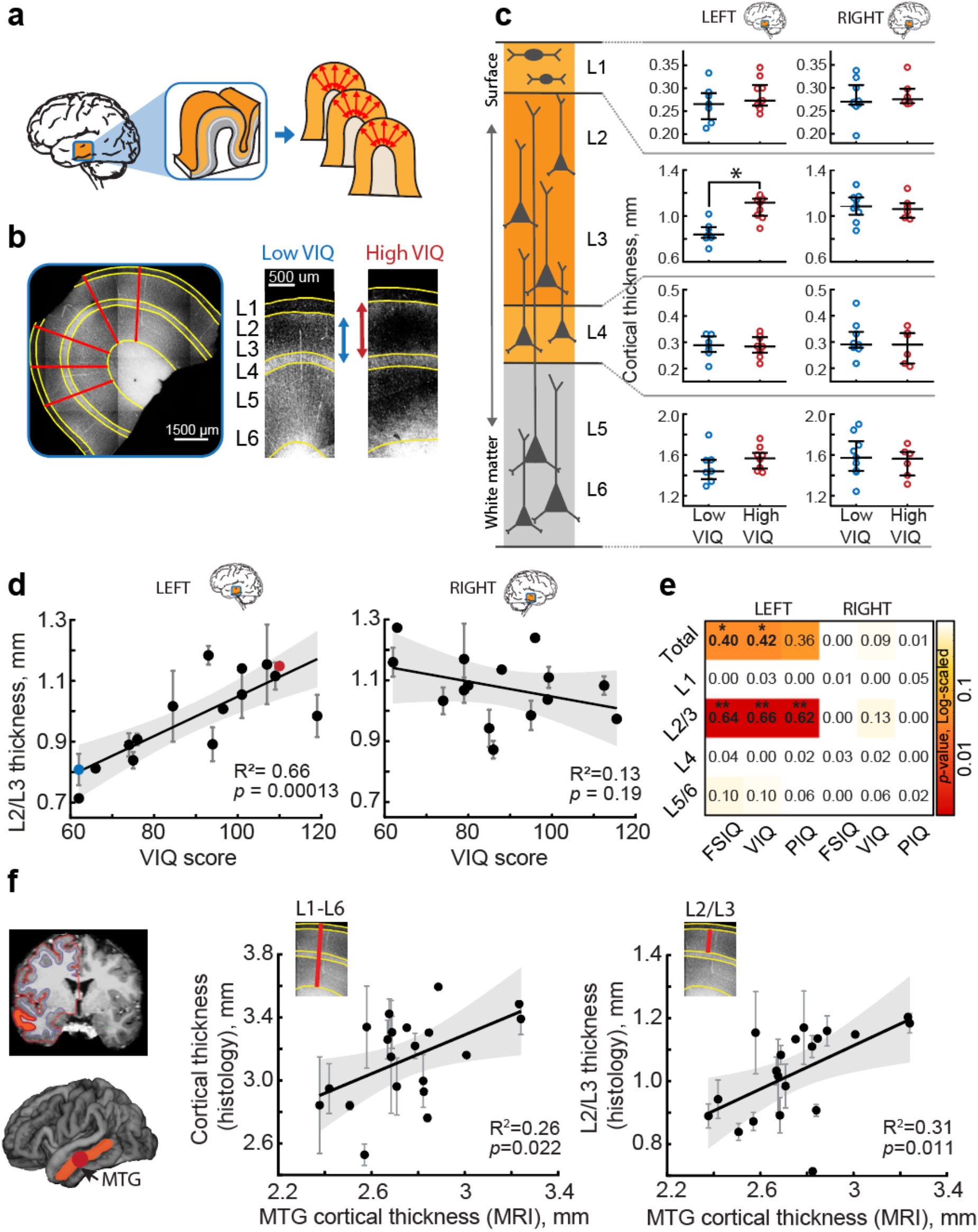
The expansion of cortical L2/L3 in the left MTG is associated with higher VIQ scores. **a** Slices from temporal lobe tissue resected during neurosurgery were DAPI-stained and imaged to determine the thickness of the cortical layers. **b** Example of cortical thickness measurements: the borders between the cortical layers were drawn on the images (yellow lines). For each patient, the average thickness was calculated along 4-5 radial lines (red) from several slices of gyral crown. **c** subjects with higher VIQ scores have thicker L2/L3 in the left MTG (Median(IQR) = 1.116(1.002-1.150) mm), than subjects with low VIQ (0.839(0.809-0.903) mm, Mann-Whitney U test, U = 32, p = 0.0021). Open circles represent the average thickness of the different cortical layers from each subject (from top to bottom: L1; L2/L3; L4; L5/L6), red for subjects with VIQ>90, blue for subjects with VIQ<90. Separately for left (left panel) and right (right panel) hemisphere (Left: low VIQ: n subjects = 7, n slices = 13, high VIQ: n subjects = 9, n slices = 22. Right: low VIQ: n subjects = 9, n slices = 25, high VIQ: n subjects = 6, n slices = 16). Here and further: black horizontal lines are median values; vertical lines are interquartile ranges. **d** L2/L3 cortical thickness positively correlates with VIQ in the left (n subjects = 16, n slices = 35, F(1,14) = 27.1), but not in the right MTG (n subjects = 15, n slices = 41, F(1,13) = 1.87). Here and further: error bars indicate SEM, shaded area represents 95% confidence bounds, insets show R2 and p-value. The blue and red data points correspond to the examples shown in b. **e** heatmap showing linear regression results (R2) for all cortical layers, both hemispheres, for Full Scale (FSIQ), verbal (VIQ) and performance (PIQ) IQ test scores. P-values are color-coded, *p < .05; **p < .01. **f** Cortical thickness in MTG quantified from MRI scans, correlates with the MTG thickness from histological quantifications shown in c and d. Left panel shows an example of an MRI scan with white-gray matter boundaries highlighted with colored lines, MTG is marked orange, the resected area is marked red. MTG cortical thickness from MRI scans of the resected MTG positively correlates with total cortical thickness (middle, n subjects = 20, n slices = 56, F(1,18) = 6.26) and L2/L3 thickness (right, n subjects = 20, n slices = 56, F(1,18) = 8.1) from histological analysis of the resected MTG.

On average, total cortical thickness, as well as L2/L3 thickness, was larger in the right MTG than left MTG. This observed asymmetry reflects a natural asymmetry in cortical thickness between left and right hemispheres, as evidenced by a recent large analysis of MRI scans of >17,000 healthy individuals from 99 datasets, where middle temporal area was shown to have higher thickness in the right hemisphere than in the left ^33^.

Intellectual performance is generally measured by FSIQ scores that are derived from both verbal (VIQ) and non-verbal, performance IQ (PIQ) scores. We therefore performed additional analysis on the relationship between L2/L3 thickness separately for VIQ, PIQ and FSIQ in the left and right MTG. The summarized linear regression results for all layers in left and right MTG, and for verbal and performance IQ scores revealed that the correlations of FSIQ, VIQ and PIQ scores with total cortical thickness are attributable to the selective expansion of L2/L3 in the left MTG. Although FSIQ, VIQ and PIQ all significantly correlated with total and L2/L3 thickness in the left MTG, variance explained (R^2^) was higher when running linear regression on VIQ than PIQ (Supplementary Fig. 2). To exclude possible confounding effects of age and gender, we ran partial correlations for the relationship between VIQ and L2/L3 thickness. We computed the zero-order and partial correlation coefficients (r) while controlling for age and gender separately for FSIQ, VIQ and PIQ and L2/L3 thickness in the left MTG. We find that the correlations remained high and significant (Supplementary Fig. 3a). Furthermore, L2/L3 thickness did not correlate significantly with age of the subjects and did not show significant difference between males and females (Supplementary Fig. 3b-c).

As cortical thickness in human subjects is usually measured using structural MRI scans, we next asked whether cortical thickness quantified using histological methods is correlated with cortical thickness quantified from MRI. To this end, we quantified cortical thickness from T1-weighted pre-surgical MRI scans using voxel-based morphometry (Fig. 1f). We selected only MTG area in the hemisphere where the resected tissue originated and calculated average MTG cortical thickness for each subject. We tested whether MRI-derived cortical thickness correlated with the histological quantifications of cortical thickness. We find that MRI and histological quantifications positively correlate with each other. Moreover, cortical thickness measured from MRI also positively correlates with the L2/L3 thickness in the gyral crown, the metric that we find most strongly related to verbal intelligence (Fig. 1f). Thus, selective expansion of the L2/L3 thickness in the left MTG strongly associates with the gain in human cognitive function, including verbal cognitive function.

### Thicker L2/L3 contains lower neuronal densities and larger cells

Next, we asked how the expansion of L2/L3 would affect the overall microstructure of these layers. We hypothesized that a thicker L2/L3 would contain larger neurons that are dispersed over a greater volume to accommodate larger dendritic arbors. To analyze the microstructure of L2/L3, we performed NeuN (neuronal nuclei) antibody staining from human frontal, temporal and parietal cortices (left and right hemisphere) in an independent group of 16 neurosurgery subjects. As L2/L3 thickness shows great variability even within the same slice of the same subject, we measured cell densities and cell size in multiple regions of interest (ROIs) covering the whole slice (24 slices, 113 ROIs). Each ROI was manually selected to include only layers 2 and 3 (Fig. 2a). Similar to previously published data ^34^, we find that neuronal density decreases from L2 to deeper L3, while the cell body area increases (Fig. 2b,c). In relation to layer thickness specifically, the neuronal density within sublayers is negatively associated with the average thickness of L2/L3: subjects with thicker L2/L3 had a less densely populated L3, and these correlations were especially large for deeper L3 (Fig 2 d,e). In addition, the cell body area positively correlated with the thickness of L2/L3. These results show that the expanded L2/L3 contain similar counts of neurons in L3, while their cell bodies are larger and more dispersed over a larger volume.

**Figure 2.**
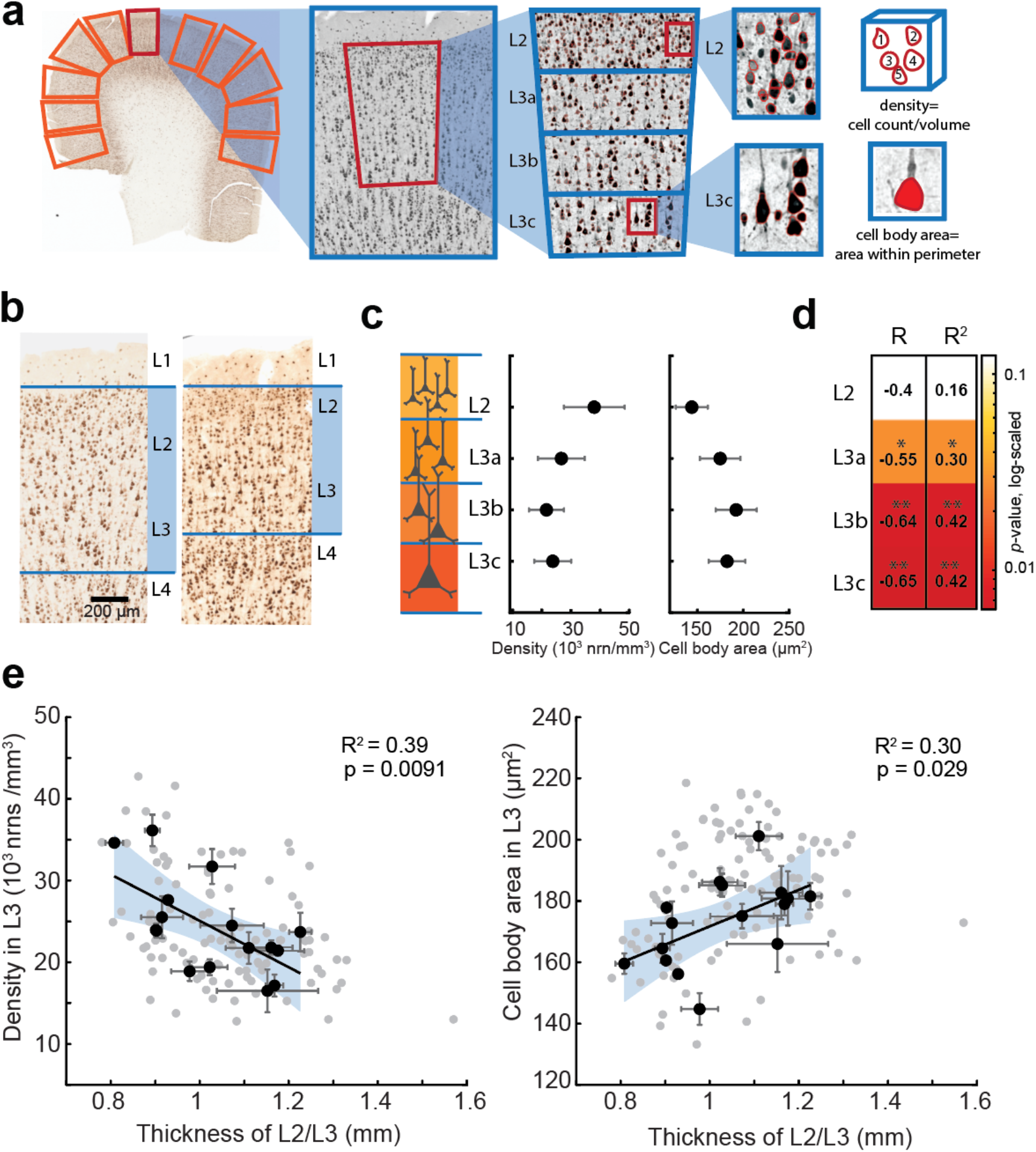
Expansion of L2/L3 in human cortex is accompanied by lower neuronal densities and larger cell body area in layer 3. **a** microstructure analysis workflow: in NeuN stained human cortical slice multiple ROIs were selected for analysis; within each ROI a region of interest was defined that included only L2/L3 and was divided in 4 sublayers of equal thickness (each at 25% of total L2/L3 thickness). The neurons were detected from the images using custom-made image-processing scripts (detected neurons are shown in red). **b** examples of NeuN stained slices from 2 subjects showing different L2/L3 thickness. **c** neuronal density decreases, and cell body area increases from L2 to deeper sublayers of L3 (black circles are mean data from 16 subjects; 24 slices, 113 ROIs). **d** results of neuronal density correlation to L2/L3 thickness per sublayer: neuronal density correlates stronger to L2/L3 thickness in deeper layer 3: correlation coefficients (R) and variance explained (R2) are shown per sublayer, p-values are color coded (*p-value<0.05; **p-value<0.01). **e** thicker L2/3 shows negative association with neuronal density in L3 (F(1,14) = 9.15) and positive association with cell body area (F(1,14) = 5.88). black circles are means per subject, n=16, gray circles are ROIs, n=113, black lines are linear regression fits to subject level data, shaded area (blue) represents 95% confidence bounds.

### Subjects with higher VIQ have larger neurons with more complex dendrites

Pyramidal cells are the principal computational units of the cortex and integrate information on their large dendrites spanning multiple layers ^12,34^. We asked whether the thicker L2/L3 layers in the left MTG from the subjects with higher VIQ also contain larger pyramidal neurons, as our data on multiple cortical brain areas in Fig. 2 suggest. To this end, we quantified the cell body diameter from the biocytin filled neurons from L2/L3 in the left and right MTG slices as shown in Figure 1. The neurons from the left MTG in subjects with VIQ >90 had significantly larger cell bodies than those from subjects with lower VIQ, while in the right MTG the distributions of cell body diameters are similar between subjects with lower and higher VIQ scores. Thus, higher VIQ associates not only with the thicker L2/L3 (Fig.1), but these layers also contain pyramidal neurons with larger cell bodies in the left MTG (Fig. 3a).

**Figure 3.**
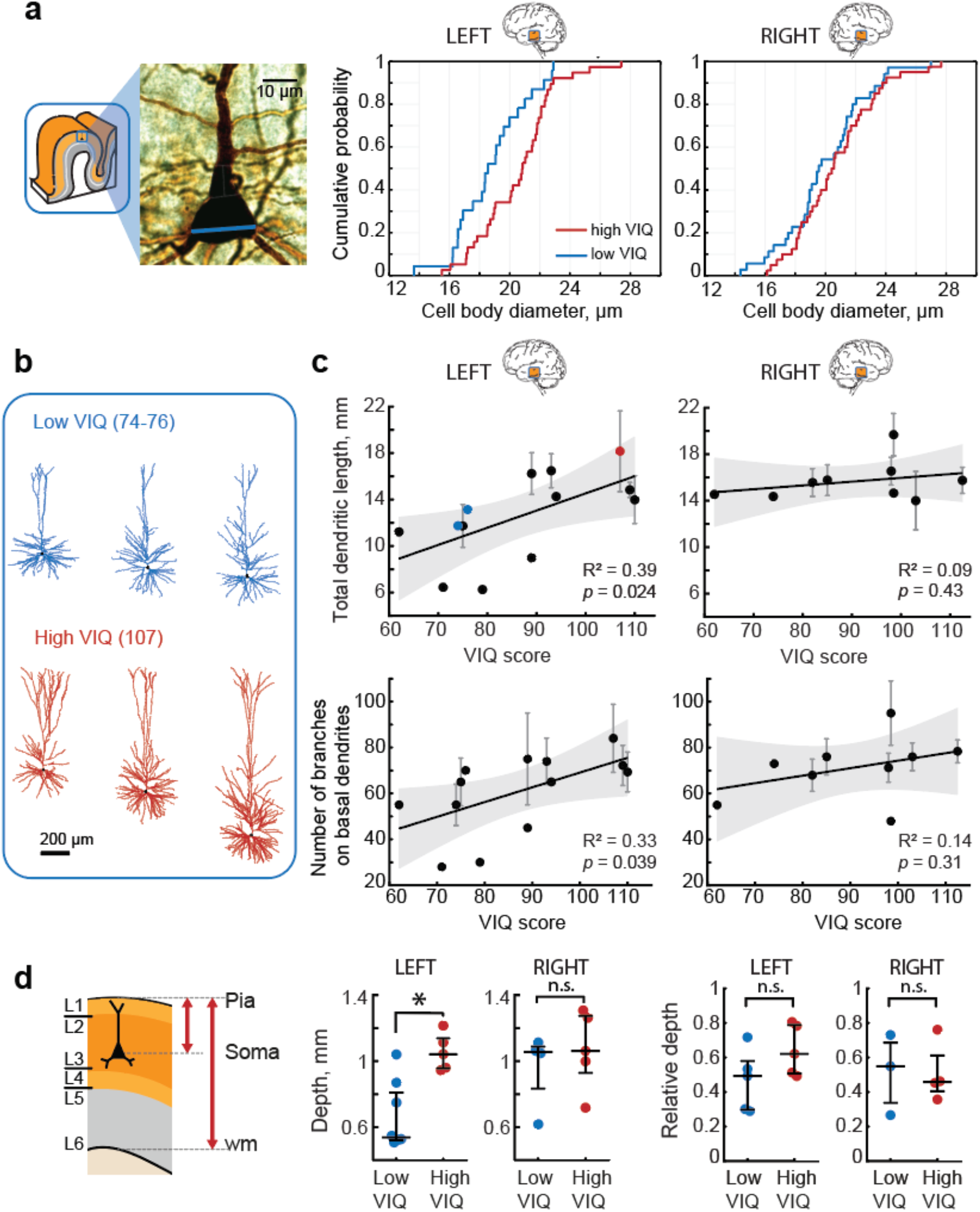
Total dendritic length (TDL) of pyramidal cells L2/L3 from the left, but not the right MTG associates with higher VIQ scores. **a** subjects with higher VIQ had larger pyramidal neurons in L2/L3: example image of a biocytin stained pyramidal neuron with soma diameter marked with blue line (left), the cumulative distribution function for all pyramidal neuron diameters in subjects with high VIQ and low VIQ for left (middle panel, n cells low VIQ=23, n cells high VIQ= 38, Mann-Whitney U test: U=268, p=0.011) and right MTG (right panel, n cells low VIQ=35, n cells high VIQ= 40, U=606, p=0.32). **b** Examples of fully reconstructed pyramidal neuronal morphologies (left MTG, L2/L3) from two subjects with low VIQ and one subject with high VIQ scores. **c** VIQ scores positively correlated with TDL (upper panel) and the number of branches (lower panel) on basal dendrites from pyramidal neurons in L2/L3 in the left (n subjects = 13, n cells = 33, TDL: F(1,11) = 6.89, number of branches: F(1,11) = 5.51), but not in the right MTG (n subjects = 9, n cells = 30, TDL: F(1,7) = 0.7, number of branches: F(1,7) = 1.19). Error bars indicate SEM, shaded area (gray) represents 95% confidence bounds. The blue and red data points correspond to the examples displayed in a. **d** Cells from both groups were recorded at similar relative depths when correcting for cortical thickness Left panel: schematic showing depth of the cell as the distance from pia to soma. Relative depth was calculated as depth divided by distance from pia to white matter. Middle panel: only in the left MTG neurons recorded from subjects with higher VIQ scores were located deeper in the cortex than those from subjects with lower VIQ (Left: low VIQ Median (IQR)= 0.537(0.523-0.810) mm, n subjects = 8, n cells = 13, high VIQ = 1.042(0.957-1.139) mm, n subjects = 5, n cells = 20, U = 38, p = 0.006. Right: low VIQ = 1.056(0.833-1.088) mm, n subjects = 4, n cells = 11, high VIQ = 1.062(0.929-1.274) mm, n subjects = 5, n cells = 19, Mann-Whitney U test: U = 18, p = 0.73). Right panel: Relative depths of the recorded cells was not different (Left: low VIQ = 0.49(0.30-0.58), n subjects = 5, n cells = 9, high VIQ = 0.62(0.51-0.79), n subjects = 5, n cells = 15, U = 21, p = 0.22. Right: low VIQ = 0.55(0.34-0.69), n subjects = 3, n cells = 8, high VIQ = 0.46(0.41-0.61), n subjects = 4, n cells = 11, U = 12, p = 1). wm: white matter.

We have previously shown that dendritic length and complexity of pyramidal cells positively associates with Full Scale IQ scores ^8^. However, it is not known whether left-lateralization of verbal function also applies to the cellular level. To test whether the total dendritic length (TDL) of pyramidal neurons in L2/L3 from the left MTG correlates with VIQ scores, we selected neurons from Figure 3a with complete dendritic trees and fully reconstructed pyramidal morphologies from the left and right MTG. In line with our findings on layer thickness and cell body size, we observed a significant positive correlation between VIQ scores and TDL for pyramidal neurons in L2/L3 in the left and not right MTG from these subjects (Fig. 3c). Moreover, VIQ scores also correlated with the number of branches on basal dendrites in these neurons (Fig. 3c), indicating that the larger dendrites in the subjects with higher VIQ are also more complex and the observed correlation with TDL is not only due to a longer apical shaft in deeper lying neurons. After controlling for age and gender, TDL also correlated with FSIQ, but not PIQ (Supplementary Fig. 3a), showing that neuronal structure in the left MTG is specifically associated with verbal cognition. Furthermore, TDL did not correlate significantly with age of the subjects and was not different between male and female subjects (Supplementary Fig. 3b-c).

Dendritic length is closely related to the location of the cell soma within cortex and deeper lying cells generally have larger dendrites ^5,35^. As subjects with higher VIQ have thicker total cortex, their pyramidal cells are on average located deeper than those from subjects with lower VIQ (Fig. 3d) and individual VIQ scores correlated with neuronal depth (Supplementary Fig. 4). To ensure that the difference in TDL was not simply the result of recording from deeper neurons in high VIQ patients, we corrected neuron depth for total cortical thickness. The relative depth of these cells within the layer was not different (Low VIQ Median(IQR) = 0.49(0.30-0.58), High VIQ = 0.62(0.51-0.79), p = 0.22) (Fig. 3d right panel) and did not correlate with VIQ scores (R^2^=0.31, p=0.097, Supplementary Fig. 4), indicating that the cells from both groups were recorded at similar relative depths within the layers. Thus, our results show that the expansion of the L2/L3 in the left MTG is accompanied by the elongation of pyramidal cell dendrites in these layers and both associate with human verbal intelligence. Longer and more complex dendrites may endow the pyramidal cells with a larger dendritic surface for forming synaptic connections, and allow separate branches of the dendritic tree to act as independent computational compartments increasing the complexity of information processing ^10,12^.

### Neurons from subjects with higher VIQ maintain fast action potentials

Dendritic tree size directly influences action potential (AP) firing of pyramidal neurons ^7,8^. It speeds up AP kinetics, increases the AP onset rapidity and allows large pyramidal neurons to better time-lock AP firing to synaptic inputs ^8^. Moreover, human pyramidal neurons from subjects with higher Full Scale IQ scores are able to maintain faster rise speeds during sustained firing. As larger TDL in pyramidal neurons from L2/L3 in the left MTG supports higher VIQ, we asked whether these neurons are also able to sustain faster AP kinetics. We performed patch-clamp recordings and recorded action potentials at different instantaneous frequencies and quantified their rise speeds (Fig 4a). We observed similar lateralization towards the left MTG: pyramidal cells in L2/L3 in the left MTG from subjects with higher VIQ scores fired APs with faster rise speeds at 21-40 Hz than those of subjects with lower VIQ. We did not observe any significant differences between VIQ groups in AP rise speeds of neurons from the right MTG (Fig. 4b,c). Finally, average AP rise speeds at 21-40 Hz positively correlated with VIQ in the left, but not in the right MTG (Fig. 4d). We checked whether one high value of AP rise speed (marked red in Fig 4d) biased the results. After excluding this data point from the regression analysis, the correlation remained strong and significant (R^2^=0.67, p=0.007). After controlling for age and gender, AP rise speeds also correlated with FSIQ, but not PIQ (Supplementary Fig. 3a). Furthermore, AP rise speeds did not correlate significantly with age of the subjects and were not different between male and female subjects (Supplementary Fig. 3b-c). Thus, subjects with higher VIQ (and FSIQ) have larger pyramidal neurons in L2/L3 of the left MTG that are able to sustain fast action potential (AP) rise speed during high frequency firing. These findings are in line with our expectation, since AP kinetics are directly influenced by dendritic morphology, where larger dendrites lead to faster AP onsets ^7,36^.

**Figure 4.**
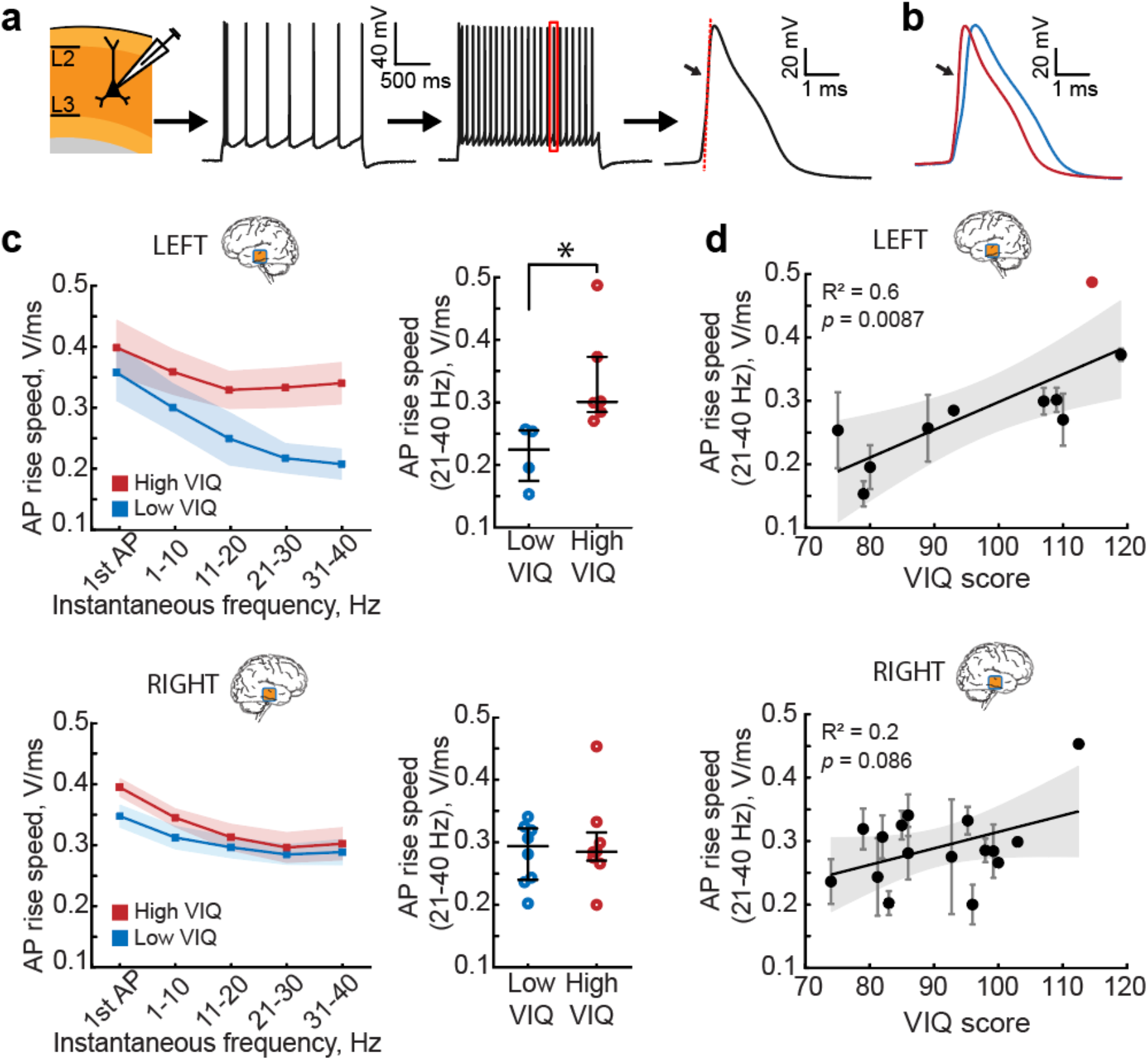
Left MTG pyramidal neurons in L2/L3 from subjects with higher VIQ scores are able to sustain fast action potential (AP) rise speed during high frequency firing. **a** Depolarizing current steps were injected in pyramidal neurons in L2/L3. APs were sorted based on instantaneous firing frequency. AP rise speed was defined as the maximum speed of the rising phase of the AP (red dotted line). **b** Example AP traces at 30 Hz from two subjects with low and high VIQ. **c** At higher frequencies, the AP rise speed is faster in neurons from subjects with higher VIQ (red), and slower in subjects with lower VIQ (blue) only in the left MTG. Shaded area represents SEM. Each data point represents an average of APs from several per subject (Left MTG, Low VIQ Median (IQR) = 224.61(174.41-255.27) mV/ms, n subjects = 4, n cells = 15, High VIQ = 300.52(284.87-372.79) mV/ms, n subjects = 6, n cells = 16 Mann Whitney U test: U = 10, p = 0.0095. Right MTG, Low VIQ = 293.79(239.81-322.17) mV/msn subjects = 8, n cells = 36, High VIQ = 284.80(270.60-315.65) mV/ms, n subjects = 8, n cells = 29, U = 67, p = 0.96). **d** AP rise speeds at higher frequencies correlate with VIQ scores only in the left MTG (n subjects =10, n cells = 31, F(1,8) = 11.9), but not in the right MTG (n subjects =16, n cells = 65, F(1,14) = 3.42). After removal of the outlier (marked in red) from the analysis, the correlation remained significant (R2= 0.67, p=0.007, F(1,7) = 14.3).

## Discussion

Our findings point to a crucial role of supragranular layers and their pyramidal cells in human verbal cognition. These results are the first to link cortical micro-organization and cellular properties in the left MTG to a specific cognitive function this area performs – verbal intelligence. Cortical thickness is one of the most robust neurobiological correlates of human intelligence, and especially the left temporal cortex in particular shows the highest correlations to general and verbal intelligence ^14,15,37–42^. On the contrary, dementia and cognitive impairment are accompanied by cortical thinning in larger areas of frontal and temporal lobes ^43,44^. Our findings demonstrate that the overall increase in cortical thickness observed in subjects with higher intelligence can be specifically attributed to the expansion of cortical L2/L3. These upper cortical layers are relatively more expanded in humans compared to other species ^1^. They are generated late in neurogenesis, and are favored by the longer period of human neurogenesis that adds novel neurons to the cortex ^45,46^. The increase in cortical surface area in humans may reflect the evolutionary shift towards increased numbers of intermediate progenitor cells ^47^. Such intermediate cell division would produce large numbers of neurons of the same subtype in the superficial layers and lead to warping of the cortical surface and gyration of the human cortex ^47^. In line with this, single-cell RNA sequencing of human MTG shows that most cells in L2/L3 surprisingly map to few transcriptomic excitatory types and are largely dominated by one type – EXC L2-3 LINC00607 FREM3 ^29^, thus supporting the role of this cell type as a principal computational unit generated by prolonged intermediate progenitor cell division in these layers. Our finding that L2/L3 expansion correlates to increased VIQ highlights that these evolutionary late additions may support better performance in verbal cognition.

The L2/L3 pyramidal cells receive rich cortico-cortical projections and have many re-excitatory connections ^48^ that they collect on their vast dendrites. In rodents they were shown to pre-amplify information ^49^ in motor cortex and induce gain modulation of deeper layer inputs in sensory cortex of rodents. This amplifying role might be even more important in multi-modal association cortices where multiple inputs have to be integrated. Deep L3 neurons have a unique pattern of dendritic maturation and have the most protracted period of developmental plasticity ^50^. They seem to be especially vulnerable and deteriorate in Alzheimer disease possibly because of their suggested role in cortico-cortical projections ^44^. Our findings corroborate the idea that their loss may substantially diminish the effectiveness of the distributed processing capacity of the neocortex.

Our results suggest that human higher cognitive functions are not only associated with a larger number of computational units, but rather with an increase in size and complexity of the individual components, the neurons. We show that individuals with higher verbal IQ scores have cells with more elaborate dendrites. This might endow pyramidal neurons with several advantages for fast and efficient computation. Firstly, large dendrites can physically contain more synaptic contacts and process more information. Secondly, separate branches of the dendritic tree can act as independent computational compartments and increase the complexity of information processing ^10,12^. These cells would need a larger volume to accommodate their vast dendrites, and thus spread more as the cortical volume expands leading to lower neuronal densities. Indeed, lower values of dendritic density were found to be associated with higher intelligence in healthy individuals ^51^. Across the brain regions, a gradient of dendritic complexity follows the function of cortical areas ^2,34,52^. Primary sensory cortices tend to have lower dendritic complexity, whereas areas involved in high-level integrative processes, contain neurons with larger and more complex dendrites in L2/L3 ^2,34,52^. As human language is built from complex hierarchically arranged structures ^53^ such as sentences, phrases, and words, it was suggested that the tree-like language structures are reflected in the tree-like structure of brain’s computational units – neurons and their dendrites ^54^.

In addition to longer dendrites, changes in the signaling speed of individual neurons must add up to increase the overall computational power of the brain. Fast signaling in the brain is a crucial requirement for efficient processing of information, and distributed brain areas of intelligence need fast signaling to coordinate their activity. Indeed, IQ scores in large groups of subjects robustly correlate with reaction times even in very simple tasks, suggesting an underlying mental speed factor ^55^. Our results provide a cellular explanation for such mental speed: we show that neurons of individuals with higher verbal IQ are able to sustain action potentials with more stable fast rise times. One of the computational consequence of the ability of neurons to maintain fast onsets of action potentials is the gain in synaptic resolution^4^. In the brain, neurons are constantly bombarded by multiple incoming synaptic inputs that they need to process, filter or pass on. The ability of the neuron to resolve and react to fast-changing synaptic inputs depends on how fast it can generate APs ^8^. Thus, faster AP kinetics and better time-locking of AP output to synaptic inputs allow large pyramidal neurons to more rapidly integrate, process and convey larger amounts of synaptic information.

The firing rates of human neurons observed in vivo indicate that maintaining fast action potentials might be especially relevant during cognitive tasks. Extracellular recording in single neurons in temporal cortex during awake craniotomy for epilepsy typically show very low baseline firing rates of <1 Hz. However, during verbal tasks, such as word pair associations learning, these neurons show task-specific shifts in their levels of activity that can last for minutes (average increase of 6-9 Hz) and can increase their activity to 30-50 Hz during specific behavioral tasks ^25,26,56–58^ These recordings were done in the same cortical area as in our study and highlight the relevance and validity of sustained high frequency firing of MTG neurons during cognitive tasks. The ability of neurons to maintain fast kinetics of APs during such sustained firing can give them an advantage in tracking fast changing synaptic inputs.

Our dataset is a unique combination of cognitive scores, structural MRI, histological assessment and single neuron measurements, however, our method has several limitations. The cortical tissue comes from subjects who undergo neurosurgery as part of their treatment for intractable epilepsy or brain tumor. Although we only use access tissue that is not part of the epileptic focus or tumor, we cannot completely rule out the influence of the disease or medication on verbal intelligence and cellular parameters. The VIQ scores from subjects in our sample are on average 90 (range 62-119), which is substantially lower than in healthy population (100). However, we do not find any correlation of VIQ scores with any of the disease history parameters, such as frequency of seizures, disease duration, and disease onset (Supplementary table 2). Furthermore, to make sure the tissue is not pathological we checked the integrity and layer structure after histological staining and controlled single cell recordings for epileptiform activity. In addition, we correlated all parameters to patients’ disease history (frequency of seizures, disease duration, and disease onset, Supplementary table 2) and found no correlations with the parameters reported in this study. Finally, we and others repeatedly demonstrated that using access tissue samples, we are able to study non-pathological properties of human circuits ^4,6,8,27,30,32,59–61^.

Another limitation of our study is that all specific cognitive test results are highly correlated and to a large degree measure the underlying common factor, general intelligence ^15^. This makes it difficult to disambiguate the unique effects of verbal and non-verbal cognitive functions. As intellectual performance is measured by FSIQ scores that are derived from verbal (VIQ) and performance IQ (PIQ), all associations of cortical architecture with VIQ inevitably will also be present with FSIQ. In our data both PIQ and VIQ highly correlate with FSIQ (correlation coefficient of FSIQ to VIQ is .907 and of FSIQ to PIQ is .902). Furthermore, performance and verbal IQ are also interrelated (in our data set correlation coefficient of VIQ to PIQ is 0.698) and share a large proportion of total variance. Thus, all associations we find of cortical parameters with verbal intelligence are similar to associations with general intelligence (measured by FSIQ).

We emphasize the association of cortical and neuronal properties to verbal intelligence primarily based on the well-documented function of the left MTG in verbal cognition. The dominant role of the left temporal lobe in verbal ability is supported by a large body of evidence from lesion studies, semantic dementia patients, findings from studies of commissurotomy patients and cortical mapping during awake neurosurgery ^19,20,62,63^. Lesions at specific locations in the left temporal lobe selectively impair specific categories of verbal reasoning such as naming of objects, tools, living things and personal names, while lesions in the right temporal lobe have much less impact on these functions ^64^. Moreover, verbal intelligence was shown to have strong structural correlates, such as cortical thickness, exclusively in the left temporal lobe, while functional correlates seem to be more symmetrically distributed ^14^. Consistent with this, our findings show that the increase in cortical thickness goes hand in hand with specific changes in superficial layers of the left temporal cortex: increased layer 2 and 3 thickness, larger dendrites, larger cell body size and decreased cell density. Furthermore, we find that variance explained (R^2^) is consistently higher for linear regressions on VIQ than on PIQ. When we control for age and gender using partial correlations, the correlation coefficients for PIQ to neuronal parameters lose their statistical significance, while VIQ and FSIQ both remain high and significant (Supplementary Fig. 3). This strengthens our conclusions that the observed effects are due to the involvement of left MTG neurons in primarily verbal aspects of intelligence.

In conclusion, our results are the first to link cortical micro-architecture of the left temporal lobe and its large and fast pyramidal neurons with verbal cognition. Our results suggest that the increased size and complexity of human neurons in these superficial layers might contribute to cortical expansion and increased human cognitive ability.

## Methods

### Human subjects and brain tissue

All procedures were approved by the Medical Ethical Committee of the VU University Medical Center, and in accordance with the declaration of Helsinki and Dutch license procedures. All subjects provided written informed consent for the use of data and tissue for scientific research. All data were anonymized.

Human cortical brain tissue was resected during neurosurgery in order to gain access to deeper pathological brain structures, typically hippocampus or amygdala. The cortical tissue originated from middle temporal gyrus (MTG, Brodmann area 21). Functional mapping was used to prevent the resection of speech areas. Subjects underwent surgery for the treatment of mesial temporal sclerosis, removal of hippocampal tumor, low grade hippocampal lesion, cavernoma, or otherwise unspecified medial temporal lobe pathology. Non-pathological cortical tissue was obtained from 59 subjects (28 males, 31 females; age 18-66 years; 26 left hemisphere and 33 right hemisphere resections). Cortical thickness was measured in 91 slices from 36 of these subjects. From 25 subjects, full morphological reconstructions of 71 neurons were obtained. Action potentials were recorded in 149 neurons from 35 subjects.

In all subjects, the resected neocortical tissue was not part of the epileptic focus or tumor and was removed to access deeper lying structures. We and others ^27–29^ have repeatedly demonstrated that using access tissue samples, one can study non-pathological properties of human circuits ^4,6,8,30–32^. We observed no correlations of cellular parameters or cognitive scores with the subject’s disease history and age (Supplementary Table 2). All anatomical, morphological and physiological data was collected and analyzed while blind to the subjects’ IQ scores.

All primary data analyses were performed blind to the cognitive tests scores of the subjects. These analyses include extraction of cortical thickness from MRI, histological staining, layer thickness quantification, cell body size and density measurements, cell body diameter measurement from biocytin stained neurons, morphological reconstructions of dendritic structure, and action potential feature extraction from electrophysiological recordings. All statistics reported in the study were performed by researchers who were not involved in the primary data analysis or IQ quantification.

### IQ scores

Full Scale IQ (FSIQ; for 58 subjects), verbal IQ (VIQ; for 50 subjects) and performance IQ (PIQ; for 51 subjects) scores were obtained using the Dutch version of the Wechsler Adult Intelligence Scale-III (WAIS-III) or WAIS-IV. The tests were typically administered within a short time before surgery as part of a neuropsychological examination. Cognitive tests were performed in the clinical setting and quantified by the clinical neuropsychologists as part of the diagnostic procedure prior to surgery. The IQ scores are calculated based on performance on the following subtests: vocabulary, similarities, information, comprehension, arithmetic, digit span, and letter-number sequencing for VIQ; picture completion, block design, matrix reasoning, digit symbol-coding, and symbol search for PIQ. For FSIQ, performance on all subtests is aggregated.

### Slice preparation

Immediately after surgical resection, the cortical tissue was transferred to carbogenated ice-cold artificial cerebrospinal fluid (aCSF) containing (in mM): 110 choline chloride; 26 NaHCO3; 10 D-glucose; 11.6 sodium ascorbate; 7 MgCl2; 3.1 sodium pyruvate; 2.5 KCl; 1.25 NaH2PO4; and 0.5 CaCL2 (300 mOsm) and transported to the laboratory. The time between resection of the tissue and the start of preparing slices was less than 15 minutes. After manual removal of the pia 350 µm-thick cortical slices were prepared in the same ice-cold solution used for transport and described above. After slicing, the slices were transferred to holding chambers filled with aCSF, containing (in mM): 125 NaCl; 3 KCl; 1.2 NaH2PO4; 1 MgSO4; 2 CaCl2; 26 NaHCO3; 10 D-glucose (300 mOsm), and bubbled with carbogen gas (95% O2/5% CO2). The slices were stored in the holding chambers for 30 minutes at 34 °C, and at least 30 minutes at room temperature prior to recordings.

### Electrophysiological recordings

Cortical slices were placed in a recording chamber with a continuous flow of oxygenated aCSF. All experiments were performed at 32-35 °C. Infrared differential interference microscopy (IR-DIC; BX51WI microscope, Olympus) was used to visualize neurons within the slices. Patch pipettes (3-5 MOhms) were filled with intracellular solution (ICS) containing (in mM): 110 K-gluconate; 10 KCl; 10 HEPES; 10 K-phosphocreatine; 4 ATP-Mg; 0.4 GTP; pH adjusted to 7.3 with KOH; 285-290 mOsm; 5 mg/ml biocytin. After establishing whole cell configuration, membrane potential responses to depolarizing current injection steps (30-50 pA step size) were recorded. Recordings were sampled at frequencies of 10 to 50 kHz and lowpass filtered at 10 to 30 kHz using Multiclamp 700A/B amplifiers (Axon Instruments). Recordings were digitized with an Axon Digidata 1440A, acquired with pClamp software (Axon) and later analyzed offline using custom-written scripts in MATLAB (R2019b, Mathworks).

### Morphological analysis

Cells were loaded with (0.5%) biocytin present in the ICS during electrophysiological recordings. Afterwards, the slices were fixed in paraformaldehyde (PFA, 4%) and the recorded cells were stained with the chromogen 3,3-diaminobenzidine tetrahydrochloride (DAB) using the avidin-biotin-peroxidase method. Next, the slices were mounted on glass microscope slides and embedded in mowiol underneath a glass coverslip. Successfully stained neurons with clear cell body contours were used for cell body diameter quantification. The cell bodies were imaged using Surveyor Software (Chromaphor, Oberhausen, Germany) with a x 100 oil objective. These images were analysed in ImageJ. The soma diameter was measured as the maximum distance from side to side at the base of the cell body, as a line perpendicular to the direction of apical dendrite. A sub-selection of these biocytin stained neurons was selected for dendritic reconstruction. This selection was based on uniform biocytin signal, presence of complete dendrites without obvious slicing artefacts, and apical dendrite reaching to layer 1. These selected neurons were digitally reconstructed using Neurolucida software (Microbrightfield) and a 100x oil objective (Olympus). After reconstruction, morphologies were checked for accurate reconstruction in x/y/z planes for presence of unconnected, missed or incompletely reconstructed dendrites. Finally, reconstructions were crosschecked by an independent researcher for false-positive/false-negative dendrites using an overlay in Adobe Illustrator between the Neurolucida reconstruction and Z-stack projection image from Surveyor Software (Chromaphor, Oberhausen, Germany), as reported previously in ^5^.

L2/L3 pyramidal neurons were identified based on morphological and electrophysiological properties, somatic depth and position within layers from DAPI stained slices. The morphological reconstructions for these cells that passed the quality control were then used to extract the total dendritic length (TDL) and number of branches on basal dendrites.

### Cortical thickness measurements

To determine cortical thickness, the fixated cortical slices that were previously recorded from, were stained with 4’,6-diamidino-2-phenylindole (DAPI) and remounted. The intensity of the fluorescence is an indication of the density of cell bodies within the cortical slice. This allowed us to differentiate the different cortical layers. Normal light and fluorescent images were taken using NeuroExplorer software (Microbrightfield). The outlines of cortical layers were tracked in ImageJ software. The cortical thickness and layer thickness were measured along radially drawn lines, parallel to fiber tracks, apical dendrites and blood vessels visible in the tissue. The lines were drawn by an independent experienced researcher to minimize errors and bias. Since no clear distinction can be made with DAPI between layers 2 and 3, and between layers 5 and 6, we treated L2/L3 and L5/L6 as single regions in the analysis. Cortical thickness varies between gyri and sulci in the cortex. Therefore, we only measured cortical thickness in the gyral crown. Cortical slices that did not clearly contain gyral crown were excluded. Cortical slices that were cut at an angle to the pia-white matter axis were identified by slicing artifacts and unclear borders between cortical layers and excluded from the analysis.

### MRI scans and cortical thickness analysis

T1-weighted brain images (1 mm thickness) were acquired with a 3T MR system (Signa HDXt, General Electric, Milwaukee, Wisconsin) as a part of pre-surgical assessment, the scans were analyzed using the Freesurfer image analysis suite (http://freesurfer.net) ^65^, previously reported in ^8^. Calculation of the cortical thickness was done as the closest distance from the grey/white boundary to the grey/CSF boundary at each vertex and was based both on intensity and continuity information from the entire three-dimensional MR volume. Neuroanatomical labels were automatically assigned to brain areas based on Destrieux cortical atlas parcellation as described in ^66^. Middle temporal gyrus was selected based on Destrieux cortical atlas parcellation in the hemisphere where the resected tissue originated. Cortical thickness at each vertex in this selected area was averaged for each subject.

### Cortical microstructure analysis from NeuN stained slices

Several slices (1-3 per subject) were fixed in paraformaldehyde (PFA, 4%) for 48 hours, transferred to phosphate buffer solution PBS + sodium azide. Slices were then cryoprotected in 30% sucrose, frozen and re-sectioned at 30 µm using a sliding microtome (Leica SM2000R). Tissue slices were stained using the Biocare Intellipath FLX slide staining automated platform. All NeuN tissue sections were pre-mounted onto gelatin coated slides, the day prior to IHC staining and first allowed to dry flat for 30-60 minutes, then were briefly rinsed in Milli-Q water. All slides were placed in 37ºC oven overnight prior to IHC staining the following day. At the day of staining, slides were peroxidase blocked in Bipocare 1X TBS wash buffer (Biocare # TWA945M), endogenous peroxidase activity was blocked using 3% hydrogen peroxidase in 1X TBS wash buffer. All slides underwent Heat Induced Epitope Retreival (HIER) methods, in 98ºC Sodium Citrate buffer, pH6.0 for 20 minutes, then allowed to cool at room temperature for 20 minutes. Next, slides were rinsed in Milli-Q water and equilibrated using 1X TBS buffer, loaded onto the Biocare intelliPath FLX® Slide Stainer and incubated on intelliPath Staining Platform using the following conditions: incubation for 10 minutes in intelliPath Background Punisher (Biocare# IP974G20), then application of 0.5µg/ml (1:2000) of NeuN mouse primary antibody (clone A60, Millipore-MAB377) in Biocare Renaissance Background Reducing Diluent (Biocare #PD950L) for 75 minutes. Next, tissue sections were rinsed in 1X TBS wash buffer and treated with BiocareMouse Secondary reagent (Mach4 kit (IPK5011 G80)) for 10 minutes, then washed in 1X TBS buffer, followed by incubation in iBiocare Universal HRP Tertiary reagent (#IPK5011 G80) for 15 minutes then rinsed in 1X TBS wash buffer. All sections were developed using a mixture of Chromogen IP FLX DAB (IPK5011 G80) and Biocare DAB Sparkle (Biocare # DS830M) applied for 1 minute. Upon autostainer run completion, all slides were unloaded into Milli-Q water, dehydrated through a series of graded alcohols, cleared in Formula 83, and coverslipped with DPX for final detection of stained neurons.

Subsequently, the images of stained subsections were acquired with 20x air on Aperio microscope at a resolution of 1 µm to 1 pixel. The quantification of cell densities and cell body sizes was performed by using custom-made MATLAB scripts (R2019b, Mathworks). Within each subsection, regions of interest (ROIs) were selected manually covering the whole slice. The border between L1 and L2 was visually identified as a characteristic sharp increase in cell body size and density, L3 to L4 border was identified at the transition from large L3 cells to 3-fold smaller L4 cell bodies. Each ROI was selected as a trapezoid with bases along the border between L1 and L2 (upper base, 500-700 µm in length), the border between L3 and L4 (lower base, 500-700 µm in lengths) and the sides parallel to the apical dendrites. The L2/L3 thickness was calculated for each ROI separately (mean length of the sides) and each ROI was split in 4 sublayers (L2, L3a, L3b and L3c defined as 25% of the L2/L3 thickness). We validated MATLAB scripts by manually quantifying neuronal parameters from ROIs from 15 slices, the manual quantification was similar to the automated quantification (average cell density quantified manually 27010 ± 3592 neurons/mm^3^, quantified with MATLAB scripts 26595 ± 4649 neurons/mm^3^).

### Statistical analysis

Statistical significance of relationships between parameters was determined using linear regression. Since multiple cells or slices were measured per subject, parameters were first averaged per subject before statistical testing. Differences between groups were tested for significance using the non-parametric two-sided Mann-Whitney U test. Corrections for multiple testing were performed according to the Benjamini-Hochberg False Discovery Rate procedure. All statistical analysis was performed using Matlab (R2019a, Mathworks). Partial correlations controlling for the effects of age and gender were computed using SPSS 26 (IBM).

## Data availability

Source data are provided with this paper. To protect the privacy of the subjects in this study, subject numbers have been randomized in Supplementary Table 1, and the source data for Supplementary Table 2 and Supplementary Fig. 3 are only available upon request from the corresponding author (NAG).

## Code availability

All customized Matlab scripts used for physiological feature extraction are available at https://github.com/INF-Rene/Morphys.

## Acknowledgements

N.A.G. received funding for this work from the Netherlands Organization for Scientific Research (NWO; VENI grant, 016.Veni.171.017). H.D.M. received funding for this work from the European Union’s Horizon 2020 Framework Programme for Research and Innovation (Human Brain Project SGA2, 785907 and HBP SGA3, 945539). Furthermore, research reported in this publication was supported by the National Institute of Mental Health under Award Number U01MH114812.

## Author contributions

Heyer DB: e-phys data acquisition, data analysis, writing manuscript

Wilbers R: data acquisition, data analysis

Galakhova A: slice histology

Hartsema Els: cell density quantification

Braak S: layer structure analysis

Hunt S: e-phys data acquisition

Verhoog MB: e-phys data acquisition

Mertens E: morphology data acquisition

Muijtjens ML:: cell density quantification

Idema S: performed neurosurgery and tissue procurement

Baayen JC: performed neurosurgery and tissue procurement

de Witt Hamer P: performed neurosurgery and tissue procurement

Klein M: IQ data acquisition

McGraw M: histology data acquisition

Lein ES: histology data acquisition

de Kock CPJ: morphology data acquisition and analysis

Mansvelder HD: study design, data analysis, writing manuscript

Goriounova NA: study design, e-phys data acquisition, histology data acquisition, data analysis, writing manuscript

## Supplementary tables

**Supplementary table 1.**
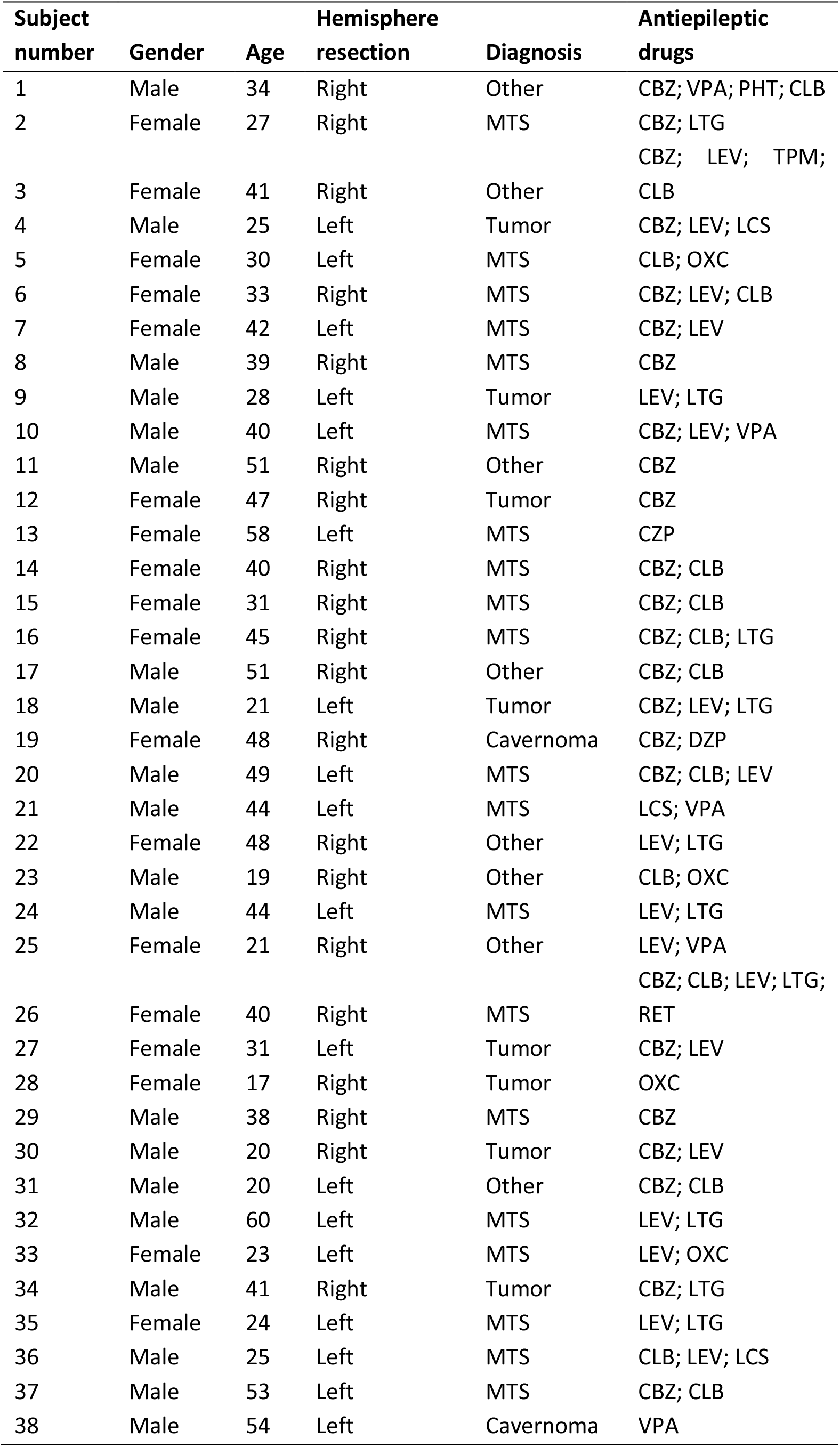
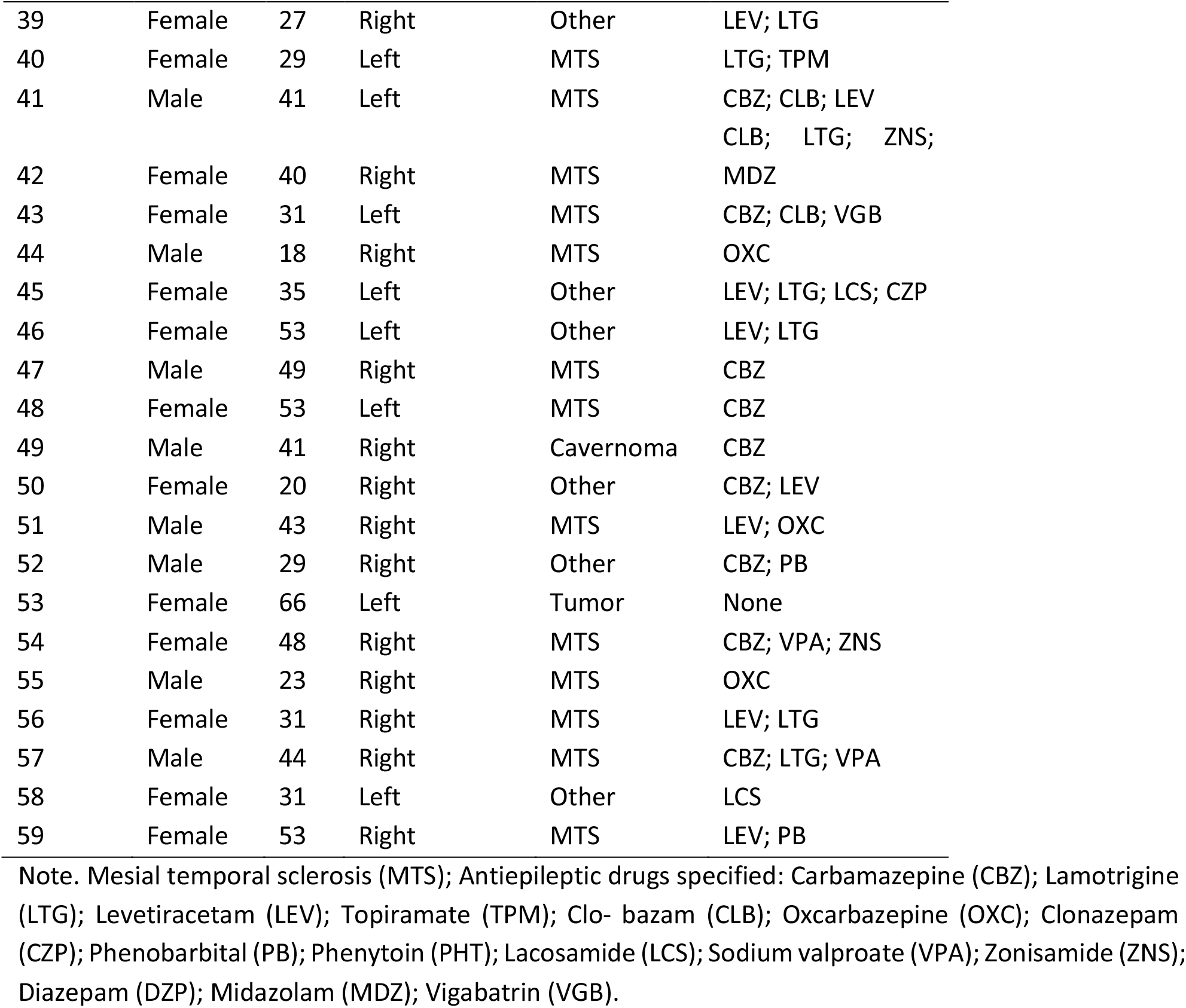
Subject details.

**Supplementary table 2.** Linear regression results indicate no relationship between the investigated parameters, disease severity or age in both left and right MTG.

| Hemisphere |  | Epilepsy onset |  | Epilepsy duration |  | Seizure frequency |  | Age |  |
| --- | --- | --- | --- | --- | --- | --- | --- | --- | --- |
|  |  | Left | Right | Left | Right | Left | Right | Left | Right |
| **VIQ** | $R^2$ | 0.039 | 0.003 | 0.020 | 0.051 | 0.166 | 0.021 | 0.005 | 0.028 |
| | $p$ | 0.66 | 0.82 | 0.70 | 0.63 | 0.33 | 0.70 | 0.82 | 0.69 |
|  | F | 1.04 | 0.09 | 0.54 | 1.45 | 5.17 | 0.56 | 0.12 | 0.78 |
|  | df | 1, 26 | 1, 27 | 1, 26 | 1, 27 | 1, 26 | 1, 26 | 1, 26 | 1, 27 |
| **Total cortical thickness** | $R^2$ | 0.129 | 0.090 | 0.222 | 0.007 | 0.052 | 0.039 | 0.013 | 0.087 |
| | $p$ | 0.58 | 0.63 | 0.33 | 0.82 | 0.66 | 0.70 | 0.82 | 0.63 |
|  | F | 2.53 | 1.49 | 4.84 | 0.11 | 0.94 | 0.57 | 0.22 | 1.43 |
|  | df | 1, 17 | 1, 15 | 1, 17 | 1, 15 | 1, 17 | 1, 14 | 1, 17 | 1, 15 |
| **L2/3 thickness** | $R^2$ | 0.148 | 0.007 | 0.080 | 0.011 | 0.110 | 0.052 | 0.021 | 0.000 |
| | $p$ | 0.58 | 0.82 | 0.63 | 0.82 | 0.63 | 0.69 | 0.76 | 0.97 |
|  | F | 2.96 | 0.11 | 1.48 | 0.17 | 2.1 | 0.77 | 0.37 | 0.002 |
|  | df | 1, 17 | 1, 15 | 1, 17 | 1, 15 | 1, 17 | 1, 14 | 1, 17 | 1, 15 |
| **TDL** | $R^2$ | 0.373 | 0.108 | 0.028 | 0.016 | 0.006 | 0.082 | 0.251 | 0.084 |
| | $p$ | 0.31 | 0.63 | 0.76 | 0.82 | 0.84 | 0.66 | 0.38 | 0.66 |
|  | F | 7.74 | 1.46 | 0.38 | 0.19 | 0.07 | 0.99 | 4.36 | 1.1 |
|  | df | 1, 13 | 1, 12 | 1, 13 | 1, 12 | 1, 11 | 1, 11 | 1, 13 | 1, 12 |
| **AP rise speed** | $R^2$ | 0.120 | 0.027 | 0.096 | 0.181 | 0.002 | 0.097 | 0.008 | 0.340 |
| | $p$ | 0.63 | 0.70 | 0.66 | 0.33 | 0.92 | 0.58 | 0.82 | 0.09 |
|  | F | 1.64 | 0.63 | 1.28 | 5.09 | 0.02 | 2.48 | 0.10 | 11.9 |
|  | df | 1, 12 | 1, 23 | 1, 12 | 1, 23 | 1, 11 | 1, 23 | 1, 12 | 1, 23 |

Note: p-values are adjusted for multiple comparisons.

## Supplementary figures

**Supplementary figure 1.**
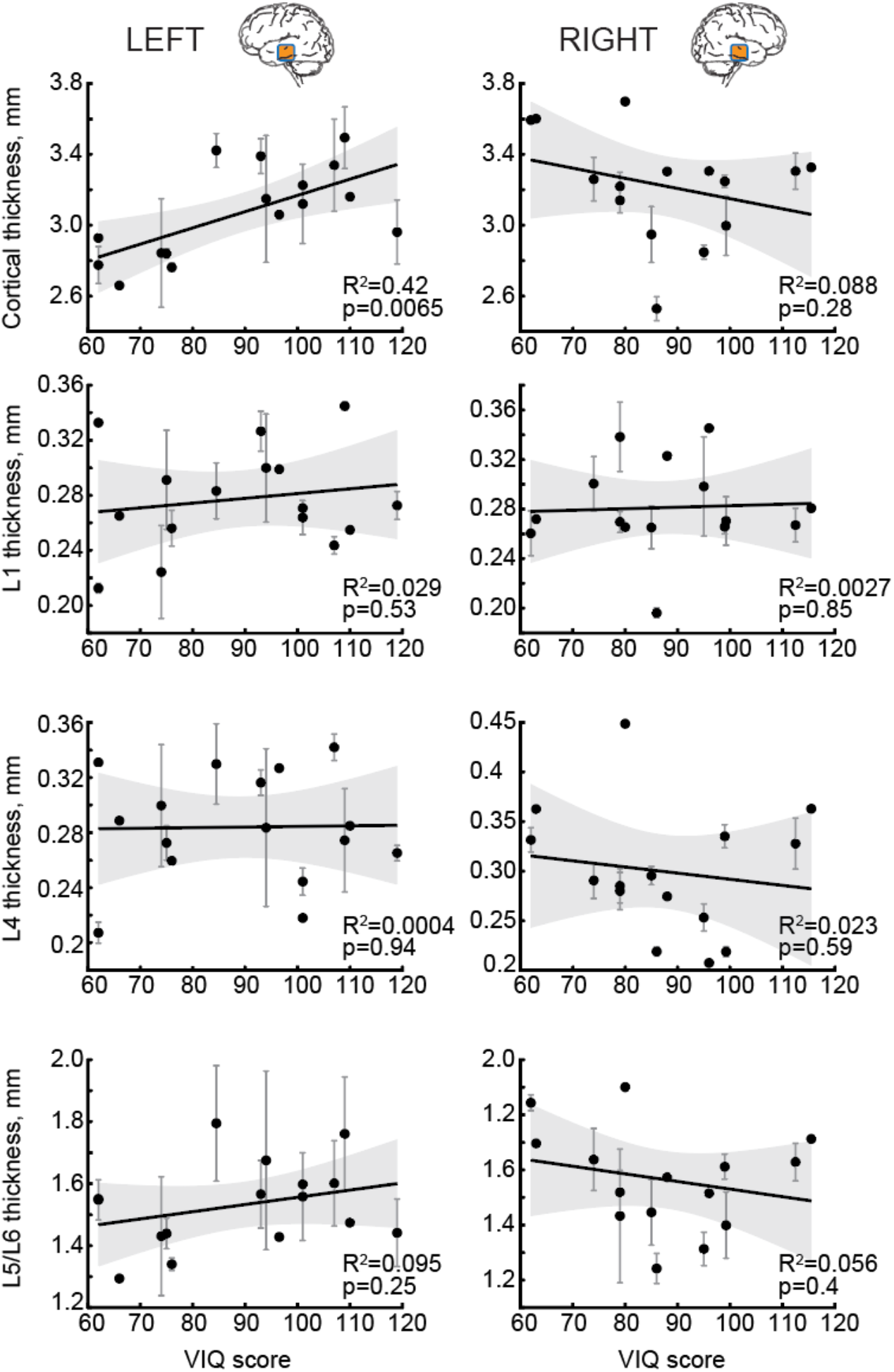
Linear regressions for VIQ with total cortical thickness and L1, L4, and L5/L6 thickness. in the left (left, n subjects = 16, n slices = 35, total: F(1,14) = 10.2, L1: F(1,14) = 0.42, L4: F(1,14) = 0.006, L5/L6: F(1,14) = 1.47) and right (right, n subjects = 15, n slices = 41, total: F(1,13) = 1.25, L1: F(1,13) = 0.04, L4: F(1,13) = 0.30, L5/L6: F(1,13) = 0.77) MTG. Only total cortical thickness in the left MTG shows significant positive correlation with VIQ.

**Supplementary figure 2.**
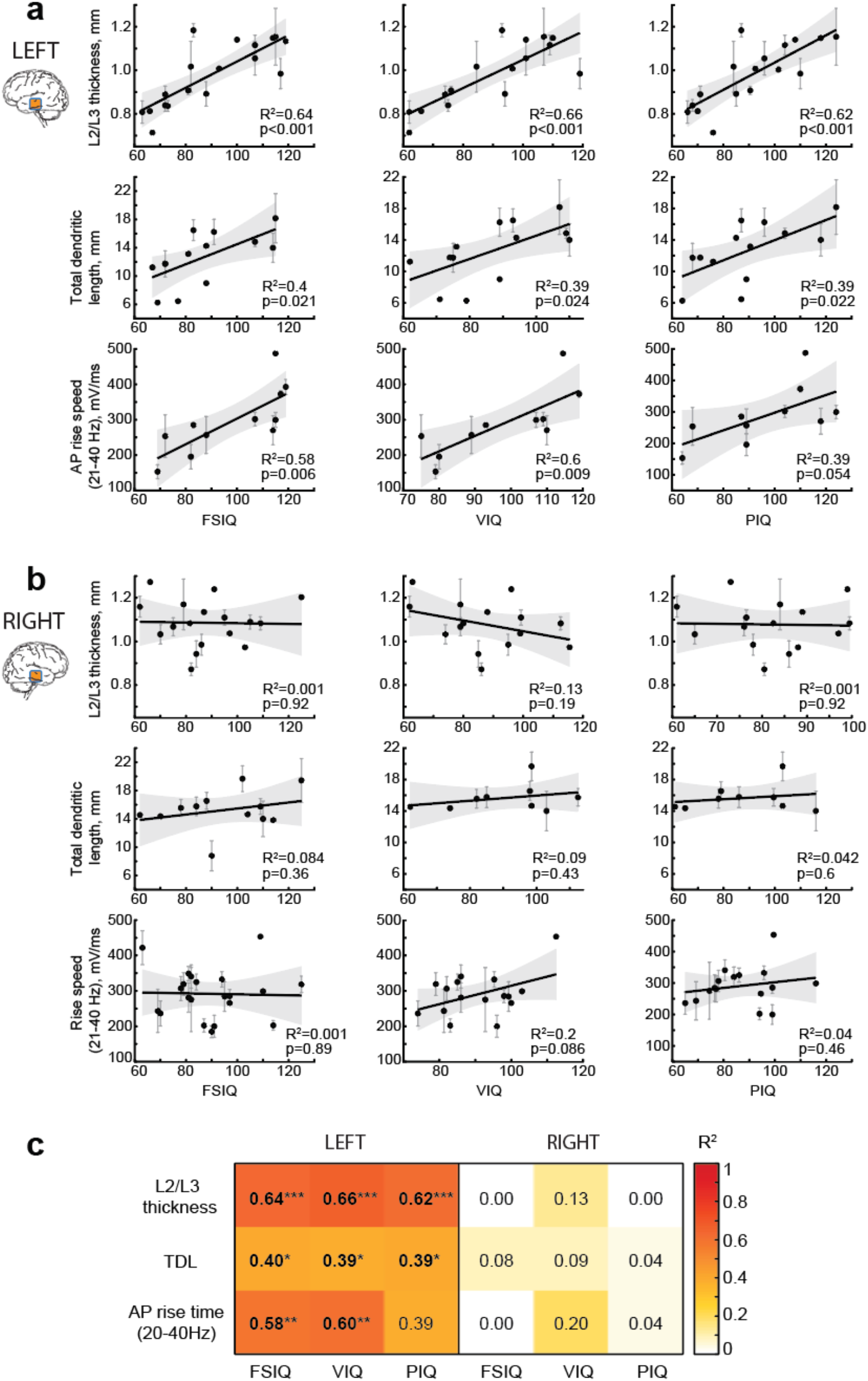
The relationship between FSIQ, VIQ and PIQ, and cortical structure and cellular properties. **a** in the left and **b** right MTG. **c** Summary of linear regression results showing R2 values for all regressions in a and b.

**Supplementary figure 3.**
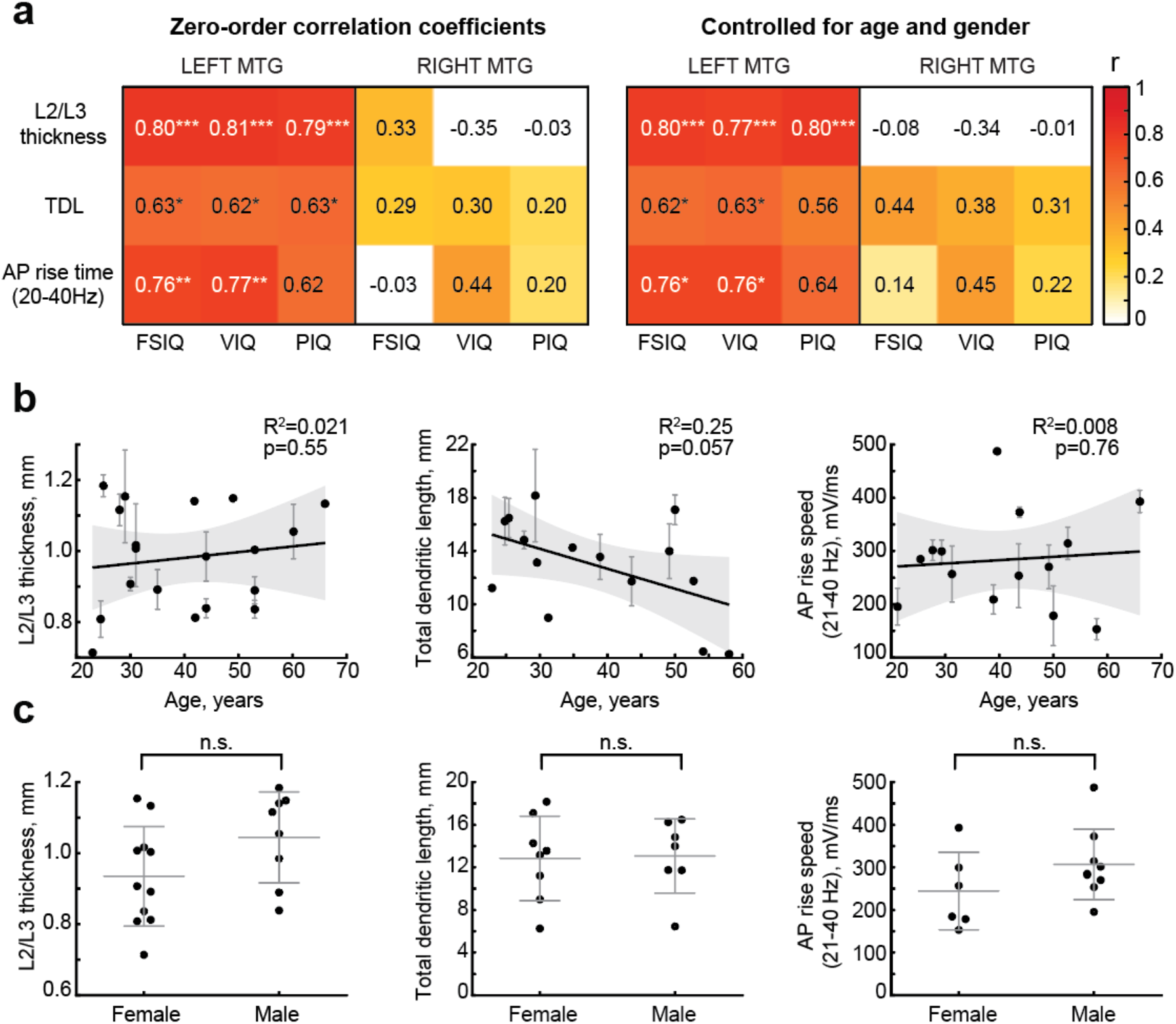
Age and gender do not influence the relationship between VIQ (and FSIQ) and cortical structure and cellular properties in the left MTG. **a** Partial correlation analysis summary showing zero-order correlation coefficients in the left panel and partial correlation coefficients controlled for age and gender in the right panel for relationship of FSIQ, VIQ and PIQ with L2/L3 thickness, total dendritic length and AP rise speed (20-40 Hz) in the left and right MTG. R remain high and significant for FSIQ and VIQ after controlling for age and gender (right). Correlation coefficients are color-coded, p-values are indicated with stars: *p < .05; **p < .01; ***<.001. **b** Age of the subjects does not correlate with L2/L3 thickness of the left MTG (n subjects = 19, n slices = 40, F(1,17) = 0.37), total dendritic length (n subjects = 15, n cells = 41, F(1,13) = 4.36) and AP rise speed (20-40 Hz, n subjects = 14, n cells = 47, F(1,12) = 0.10) of neurons in the left MTG. R2 and p-values are shown as insets. **c** Female and male subjects do not significantly differ in their L2/L3 thickness of the left MTG (females M(SD) = 0.934 (0.140) mm, males M(SD) = 1.044(0.127) mm, z(n = 19) = -1.53; p = 0.127), total dendritic length (females M(SD) = 12.84(3.96) mm, males M(SD) = 13.07(3.49) mm, z(n = 15) = -0.174; p = 0.862), and AP rise speed (20-40 Hz) of neurons in the left MTG (females M(SD) = 244.28(91.03) mV/ms, males M(SD) = 307.01(82.63) mV/ms, z(n = 15) = -1.36; p = 0.175).

**Supplementary figure 4.**
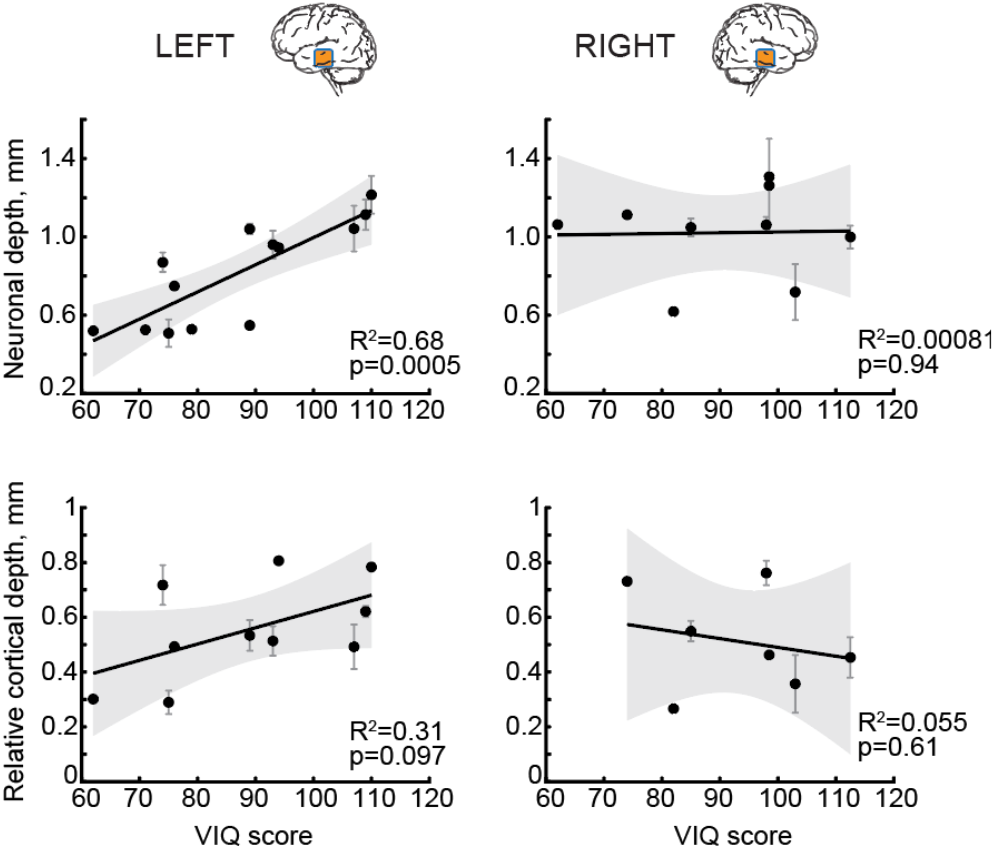
VIQ positively correlates with the absolute, but not relative cortical depth of neurons in the left MTG. Relationship between VIQ and absolute and relative cortical L2/L3 depth of neurons in the left (left, absolute: n subjects = 13, n cells = 33, F(1,11) = 23.4, relative: n subjects = 10, n cells = 24, F(1,8) = 3.54) and right (right, absolute: n subjects = 9, n cells = 30, F(1,7) = 0.006, relative: n subjects = 7, n cells = 19, F(1,5) = 0.29) MTG used for morphological analysis in Figure 3b-d.

